# Accounting for chimerism in demographic inference: reconstructing the history of common marmosets (*Callithrix jacchus*) from high-quality, whole-genome, population-level data

**DOI:** 10.1101/2025.02.11.637666

**Authors:** Vivak Soni, Cyril J. Versoza, Eric J. Vallender, Jeffrey D. Jensen, Susanne P. Pfeifer

## Abstract

As a species of considerable biomedical importance, characterizing the evolutionary genomics of the common marmoset (*Callithrix jacchus*) is of significance across multiple fields of research. However, at least two peculiarities of this species potentially preclude commonly utilized population genetic modeling and inference approaches: a high frequency of twin births and hematopoietic chimerism. We here investigate these effects within the context of demographic inference, demonstrating via simulation that neglecting these biological features results in significant mis-inference of the underlying population history. Based upon this result, we develop a novel approximate Bayesian inference approach accounting for both common twin-births as well as chimeric sampling. In addition, we present novel population genomic data from 15 individuals sequenced to high-coverage, and utilize gene-level annotations to identify neutrally evolving intergenic regions appropriate for demographic inference. Applying our developed methodology, we estimate a well-fitting population history for this species, which suggests robust ancestral and current population sizes, as well as a size reduction roughly 7,000 years ago likely associated with a shift from arboreal to savanna vegetation in north-eastern Brazil during this period.

## INTRODUCTION

Characterized by exudivorous feeding habits and small habitat ranges (∼5,000 to 65,000 m^2^), the common (or white-tufted-ear) marmoset (*Callithrix jacchus*) is a platyrrhine native to east-central Brazil (Rylands and Faria 1993; Rylands et al. 2009; Garber et al. 2019). Due to its diminutive size (∼250 g), early sexual maturity (∼15 to 18 months of age), short gestation period (∼145 days), and high fecundity (being able to give birth to up to four offspring in a single pregnancy, with twin births being the norm), this species has risen in biomedical prominence as a commonly used model for the study of both human neurodevelopment disorders (*e.g*., Miller et al. 2016; Philippens and Langermans 2021) as well as infectious disease dynamics, with the latter partly owing to their seemingly reduced major histocompatibility complex diversity relative to other mammals (Antunes et al. 1998; Wu et al. 2000; Carrion and Patterson 2012).

Together with a high frequency of twin births, *C. jacchus* is also noteworthy for the frequent observation of hematopoietic chimerism – a rare phenomenon amongst primates. As a consequence, marmoset blood samples have been shown to contain genetic material from both the sampled individual as well as from their twin sibling (Hill 1932; Wislocki 1939; Benirschke et al. 1962; Gengozian et al. 1969; Ross et al. 2007). Importantly, Ross et al. (2007) found that chimerism in marmosets was not only limited to blood samples, but was in fact identified in every sampled tissue examined. However, Sweeney et al. (2012) subsequently suggested the possibility that blood infiltration may give rise to such apparent chimerism across tissues. This result was largely confirmed by del Rosario et al. (2024), who found that chimerism present in liver, kidney and brain tissues was the result of myeloid and lymphoid cell lineages derived from hematopoietic stem cells, and that blood samples contained the greater contribution of sibling nuclei relative to other examined tissue types (see Figure 1 in del Rosario et al. 2024, and the accompanying commentary of Chiou and Snyder-Mackler 2024).

**Figure 1:**
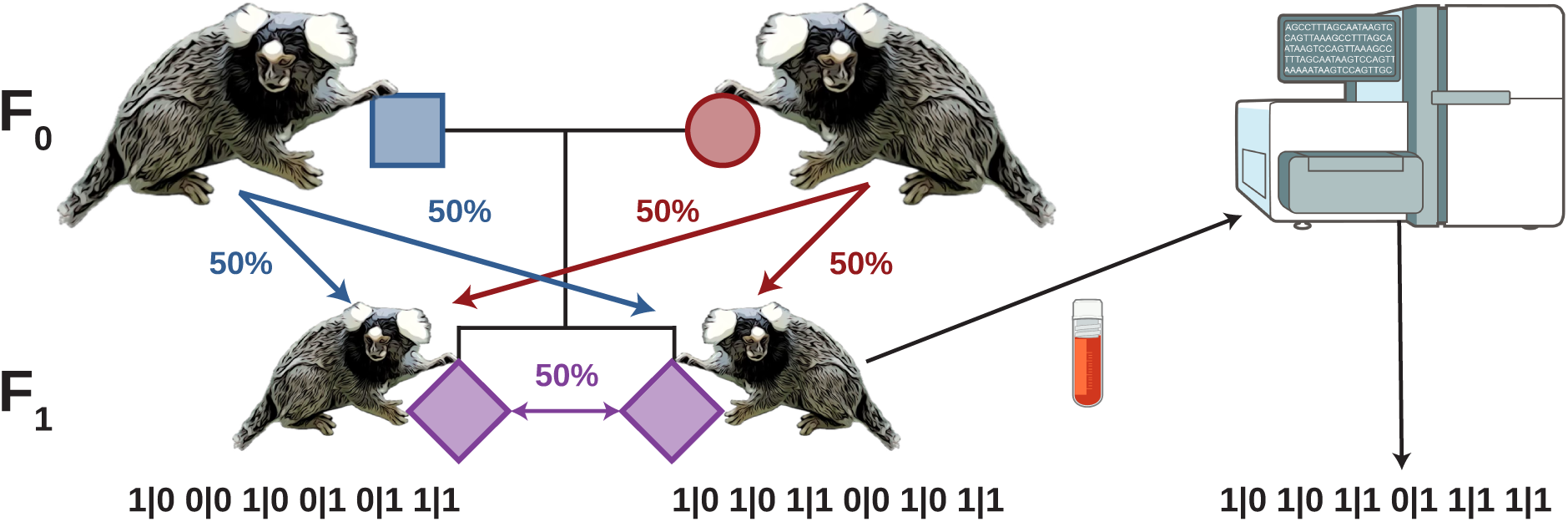
Modelling of chimerism. Marmoset blood samples contain genetic material from both the sampled individual as well as from their dizygotic (fraternal) twin (at a 1:1 ratio on average). To model chimerism, we thus combined the genotypes from an individual and its non-identical twin such that a mutation present in at least one of the twins was considered to be a SNP. For example, if one twin had a genotype of 0|1 and the other twin had a genotype of 1|1, the constructed chimeric genotype was considered to be 1|1. Marmoset cartoons were adapted from a picture taken by Eric Kilby and shared under the CC BY-SA 2.0 license; the blood vile and sequencer cartoons were taken from NIAID Visual & Medical Arts. (07/10/2024). Cryo Blood Vial and Next Gen Sequencer. NIAID NIH BIOART Source. bioart.niaid.nih.gov/bioart/87 and bioart.niaid.nih.gov/bioart/386.

Although it may be tempting to try to side-step the effects of chimerism when performing genetic studies in marmosets by focusing on single births, it is noteworthy that single births are not only rare, but also tend to themselves be chimeric owing to fetal resorption of a dizygotic twin *in utero* (Jaquish et al. 1996; Windle et al. 1999). It has additionally been proposed that one may better utilize samples from tissues potentially less impacted by chimerism (*e.g*., fingernails, as assessed from lower levels of heterozygosity; Yang et al. 2023); however, the lower (but non-zero) levels of chimerism in non-blood tissue only exacerbate the uncertainty in modelling. Hence, in terms of the sample itself, it appears difficult to reliably avoid the contribution of genetic material from the unsampled twin. Moreover, even if it were possible to sample genetic material from a true single individual in any given generation, the long-term, multi-generation genetic transmission resulting from standard twin-births would nonetheless require investigation in order to understand population-level allele frequency dynamics in this species, given this inherent violation of standard modeling assumptions.

The first marmoset genome, assembled from whole-genome shotgun plasmid, fosmid, and BAC end sequences, was published in 2014 (The Marmoset Genome Sequencing and Analysis Consortium 2014). Additional studies have since continued to improve upon this initial assembly (see the review of Vallender 2019), filling in gaps and refining gene annotations, with the most recent reference genome now exhibiting 98.3% completeness (Yang et al. 2021). Population genomic inference has thus far been limited to characterizing general levels of genetic diversity and divergence in the species (Faulkes et al. 2003; Malukiewicz et al. 2014; Yang et al. 2021; Yang et al. 2023; Mao et al. 2024), as well as performing genomic scans, some of which have implicated genes putatively involved in twinning as having experienced long-term positive selection (Harris et al. 2014). Importantly however, no study to date has attempted to model the effects of chimerism on population genetic inference, though one may readily hypothesize that effectively treating sampled chimeras as a single individual (*i.e*., as though sequenced from a single non-chimeric blood / tissue sample) may well have important implications for observed levels and patterns of genetic variation, and thus on downstream evolutionary inference. For example, Mao et al. (2024) found that a non-chimeric closely related platyrrhine (owl monkeys, for which the rate of twinning is also considered to be extremely low; Huck et al. 2014) was characterized by considerably lower divergence to humans compared with marmosets, whilst Harris et al. (2023) found that marmosets had a generally reduced relative heterozygosity. However, the potential effects of twinning and chimerism themselves on these observations were left unexplored, and it thus remains unclear whether these unusual reproductive dynamics, or, for example, fundamental differences in population size and/or mutation rates, better explain these patterns.

### Disentangling the effects of neutral and selective evolutionary processes

As a focal point of population genetic inference is on quantifying the relative roles of neutral and selective processes in governing levels and patterns of genetic variation, and as chimerism likely plays an additional role in shaping this variation, it is thus necessary to evaluate expectations associated with this inherent ‘twin-sampling’. However, chimerism and twinning aside, disentangling these competing processes is already relatively challenging (for a discussion, see the reviews of Charlesworth and Jensen 2022; Jensen 2023). As but one example, population growth, background selection (BGS; Charlesworth et al. 1993), and recurrent positive selection (Maynard Smith and Haigh 1974) can all result in a similar skew towards rare alleles when examining allele frequency distributions (*i.e*., the site frequency spectrum [SFS]; Kim 2006; Jensen et al. 2007; Nicolaisen and Desai 2012, 2013; Ewing and Jensen 2014, 2016; Johri et al. 2021; Soni et al. 2023; and see the reviews of Charlesworth and Jensen 2021, 2024). Apart from natural selection and genetic drift, heterogeneity in underlying mutation and recombination rates across the genome can also modify these expectations in significant ways (Dapper and Payseur 2018; Samuk and Noor 2022; Soni et al. 2024a). For these reasons, the case has been made that prior to evaluating genomes for evidence of relatively uncommon evolutionary processes such as positive and balancing selection, one must first construct an evolutionarily appropriate baseline model accounting for these constantly operating evolutionary processes (Bank et al. 2014; Johri et al. 2022a).

Importantly, the appropriate approach to take when accounting for the potentially confounding effects of, for example, selection inference with demographic inference, will depend on the specific details of the genome architecture of the species in question. In coding-dense genomes, the large genomic fraction of directly selected sites implies that there may be few genomic regions that are free from the effects of either direct selection or selection at linked sites. As such, the majority of genomic regions are likely shaped by both demography and selection (*e.g*., Irwin et al. 2016; Sackman et al. 2019; Jensen 2021; Morales-Arce et al. 2022; Terbot et al. 2023a,b; Howell et al. 2023; Soni et al. 2024b). Under this scenario, it is necessary to simultaneously infer population history and selection jointly, and approximate Bayesian computation (ABC) approaches have been developed for this purpose (Johri et al. 2020, 2021, 2023) – though it should be noted that, owing to the large number of underlying parameters involved, these approaches have remained limited to simplified demographic models to date.

By comparison, there are notable advantages to performing evolutionary inference in species with coding-sparse genomes (including marmosets and other primates). Owing to the prevalence of neutral sites at sufficient recombinational distances from functional sites, such that they are unlikely to experience BGS effects (Charlesworth et al. 1993), so-called ‘two-step’ inference approaches become viable (see Soni and Jensen 2025). Here, population history can be inferred from neutral intergenic regions, and numerous well-performing neutral demographic estimators have been developed for such purposes (*e.g*., Gutenkunst et al. 2009; Excoffier et al. 2013; and see the review of Beichman et al. 2018). Conditional on the population history inferred in this first step, selective processes can then be inferred using functional sites in a second step. Crucially, this baseline model accounting for population history, structure, and gene flow, together with the action of purifying and background selection in and around functional regions, has been found to be important for reducing false-positive rates when scanning for the episodic effects of positive or balancing selection (*e.g.,* Barton 1998; Przeworski 2002; Jensen et al. 2005; Poh et al. 2014; Harris and Jensen 2020; Soni and Jensen 2024; and see Johri et al. 2022b). Such multi-faceted analyses of the inter-related evolutionary processes of mutation and crossover / non-crossover rates, with population history and purifying selection, have been executed in a small number of non-chimeric primate species, including in both a haplorrhine (humans; Kong et al. 2002; DeGiorgio et al. 2016; Carlson et al. 2018; Johri et al. 2023; Ragsdale et al. 2023; Soni and Jensen 2025) as well as a strepsirrhine (aye-ayes; Versoza et al., 2024, 2025; Versoza, Lloret-Villas et al. 2024; Soni et al. 2024c; Soni, Versoza et al. 2024; Soni et al. 2025; Terbot et al. 2025).

### Incorporating chimerism into an evolutionary baseline model

Given the long-standing and compelling evidence for chimerism in marmosets (see the review of Malukiewicz et al. 2020), together with the high rate of twin births, these features will necessarily be an important component of the evolutionary baseline model for this species in order to fully characterize neutral expectations. Notably, accounting for species-specific biology is not particularly uncommon in baseline model construction. For example, in species characterized by strong reproductive progeny skew (as observed *e.g.*, in both marine and terrestrial broadcast spawners, as well as multiple pathogens), one must similarly account for the impacts of such a Wright-Fisher (WF) model violation on downstream inference, owing to related modifications of expected neutral patterns of variation (Eldon and Wakeley 2006; Matuszewski et al. 2018; Sackman et al. 2019; Sabin et al. 2022; and see the reviews of Tellier and LeMaire 2014; Irwin et al. 2016). Importantly, a neglect of this biological reality in affected organisms has been demonstrated to result in both a mis-inference of population growth as well as the false detection of widespread positive selection (*e.g*., Durrett and Schweinsberg 2004; Hallatschek 2018).

In this study, we thus uniquely construct an appropriate population genomic baseline model for *C. jacchus*, and firstly examine the effects of hematopoietic chimerism on equilibrium expectations of levels (*e.g*., *θ_w_*, Waterson 1975) and patterns (*e.g*., the SFS) of genetic diversity. Secondly, based on these observed deviations, we quantify the extent to which common demographic inference approaches may be biased by the unaccounted for presence of chimerism. Thirdly, we develop a novel ABC framework for inferring population history in the presence of both twin-births and chimerism, and present a simulation study evaluating the performance of this approach. Finally, we present novel population genomic data from 15 unrelated common marmoset individuals sequenced to high-coverage and, utilizing gene-level annotations to identify genetic variation in intergenic regions appropriate for demographic inference, we apply our ABC approach to estimate a well-fitting population history for the species. Taken together, these results demonstrate the importance of accounting for chimerism when performing demographic inference in chimeric species, with the resulting demographic model in *C. jacchus* suggesting a population bottleneck roughly 7,000 years ago, followed by a partial recovery. Paleoclimatic and palynological studies indicate a shift from arboreal to savanna vegetation in north-eastern Brazil during the timeframe of this bottleneck – an event that would be anticipated to impact a species that relies on arboreal locomotion.

## RESULTS AND DISCUSSION

### Whole genome, population-level data

In order to infer the population history of common marmosets (*C. jacchus*), we whole-genome sequenced 15 unrelated individuals previously housed at the New England Primate Research Center to mean depths of 35X per individual (Supplementary Table S1). Following best practices in the field (Pfeifer 2017), we mapped individual reads to the current marmoset reference genome and subsequently called, genotyped, and filtered variants using the Genome Analysis Toolkit workflow (van der Auwera and O’Connor 2020). Additionally, to account for the effects of both direct and background selection – two evolutionary processes previously shown to bias demographic inference (*e.g*., Johri et al. 2021; and see the reviews of Charlesworth and Jensen 2021, 2024) – we excluded any sites within, or sufficiently close to, functional regions prior to downstream analyses (see “Materials and Methods” for further details). The final dataset of putatively neutral regions consisted of 1.4 million autosomal, biallelic single nucleotide polymorphisms (SNPs) with a transition-transversion ratio of 2.11 in the accessible genome (Supplementary Table S2). Table 1 provides the means and standard deviations of *θ_w_* (Waterson 1975), Tajima’s *D* (Tajima 1989), and the number of singletons across these putatively neutral regions.

**Table 1:** Means and standard deviations (SD) of *θ_w_* (Waterson 1975), Tajima’s *D* (Tajima 1989), and the number of singletons across putatively neutral regions in the empirical data, calculated across 10kb windows.

| | $\theta_W$ | Tajima's $D$ | # singletons |
| --- | --- | --- | --- |
| **mean** | 0.0010 | 0.2548 | 0.0007 |
| **SD** | 0.0009 | 1.1484 | 0.0013 |

### Evaluating the impact of chimerism on the performance of common SFS-based estimators of population history

As marmoset blood samples have been shown to contain genetic material from both the sampled individual as well as from their dizygotic (fraternal) twin (Hill 1932; Wislocki 1939; Benirschke et al. 1962; Gengozian et al. 1969; Ross et al. 2007), we modelled chimerism by simulating a non-Wright-Fisher (non-WF) model in SLiM4 (Haller and Messer 2023) in which monogamous mating pairs produce non-identical twin offspring. The genotypes of these twins were combined post-simulation (such that a mutation present in at least one of the twins was considered a SNP) to create a chimeric individual (Figure 1; and see “Materials and Method” for further details). To evaluate the performance of commonly used SFS-based demographic inference methods, we simulated a single neutrally evolving population under four demographic scenarios: (1) population equilibrium (with a single parameter *N_current_*, the population size at the time of sampling), as well as three instantaneous population size change scenarios (with three parameters each: *N_ancestral_*, the population size prior to the size change; *T*, the time of the size change in *N_ancestral_* generations; and *N_current_*) – namely, (2) population expansion (population size doubling), (3) population contraction (population size halving), and (4) severe population contraction (population size reduced to 0.1*N_ancestral_*). These scenarios were simulated under both our chimeric model as well as a standard WF model in order to compare demographic inference power with and without chimerism. We performed demographic inference with the coalescent-based estimator fastsimcoal2 (Excoffier et al. 2013) and the diffusion approximation of *δaδi* (Gutenkunst et al. 2009), both of which are neutral estimators that fit a demographic model to the observed SFS.

Generally, chimerism resulted in mis-inference of demographic parameters across all four population histories (see Figure 2 for the results of the demographic inference of the equilibrium population, and see Supplementary Figures S1-S3 for the three population size change scenarios). Notably, the variance on parameter estimates was greatly increased under chimerism relative to the WF population, with the direction of mis-inference depending on the underlying population history. For example, the time of size change, *T*, was underestimated by both demographic estimators for the expanding population (Supplementary Figure S1) but overestimated in the contracting populations (Supplementary Figures S2 and S3). This likely owes to the fact that both chimerism and population growth are acting to reduce the correlation between underlying genealogies across the genome, and thus chimerism results in a mis-inference towards more recent growth; whereas chimerism and population decline are acting in opposite directions with regards to these genealogical correlations, and thus chimerism results in a mis-inference towards a more ancient decline. As a general trend, *θ_w_* (Waterson 1975) was reduced under chimerism relative to the WF model. This pattern is likely driven by monogamous non-identical twin births reducing the effective population size, whilst the increase in Tajima’s *D* (Tajima 1989) relative to the WF model is the result of fewer singletons in the chimeric genome (due to the combining of twin genomes), and thus a skew toward higher frequency alleles in the SFS. Combined with the increased variance in inferred parameters, these results suggest that neutral demographic estimators are not particularly well suited for inferring the population histories of chimeric species; thus, we have here developed a novel ABC approach for this purpose.

**Figure 2:**
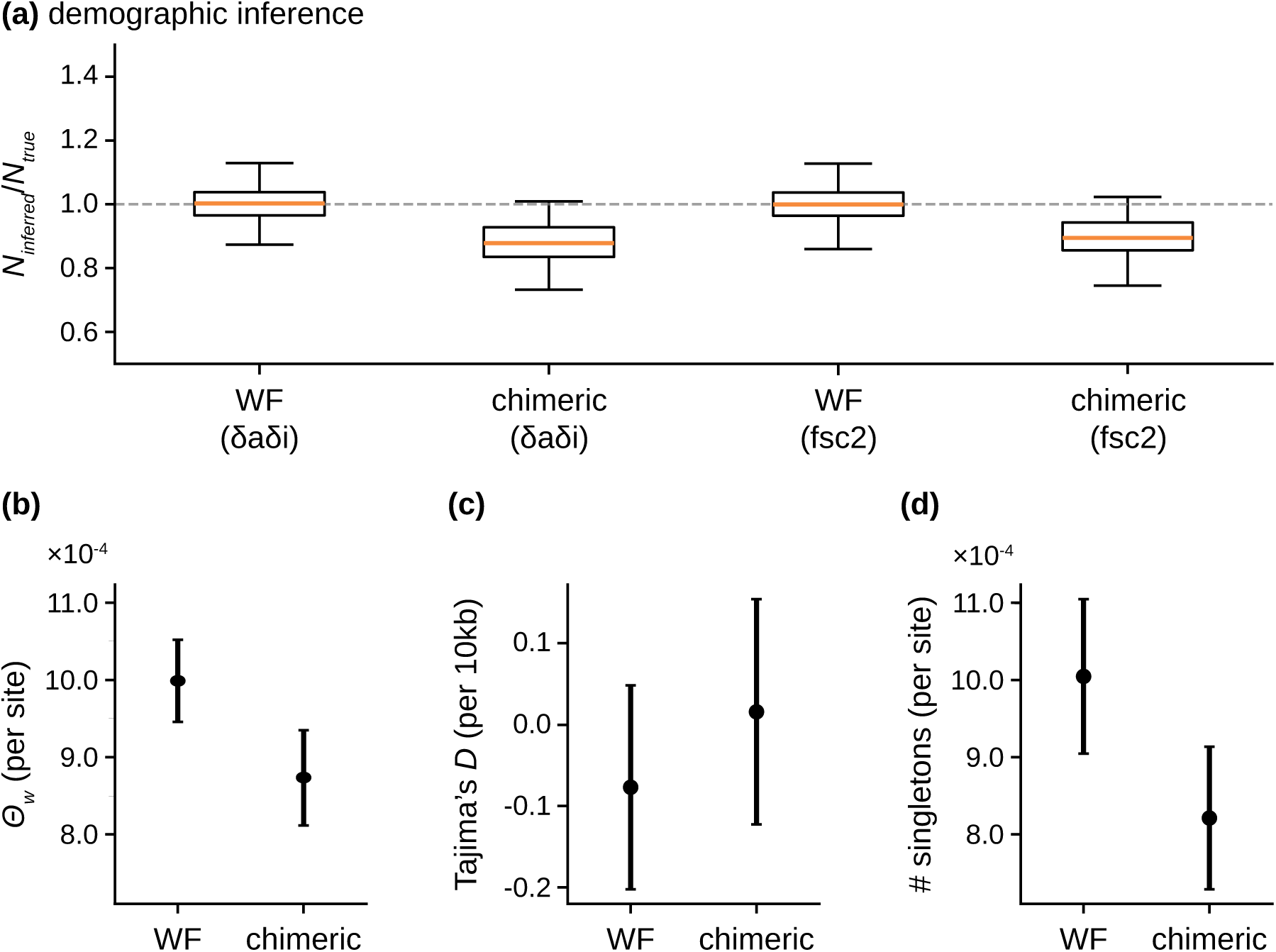
Demographic inference and summary statistics for simulations of an equilibrium, neutrally evolving population, across 100 simulation replicates, comparing Wright-Fisher (WF) and chimeric models. **a)** Demographic inference results from *δaδi* (Gutenkunst et al. 2009) and fastsimcoal2 (Excoffier et al. 2013). The y-axis represents the inferred value of the single parameter, the population size (*N*), relative to the true value of this parameter (*N_inferred_*/*N_true_*, with a value of 1 indicating that the inferred and true values are in agreement). The orange line represents the mean inference value across 100 simulation replicates, with boxes representing the 25 and 75 percentiles, and whiskers representing minimum and maximum values. **b-d)** Summary statistics including Watterson’s *θ* (*θ_w_*; Waterson 1975) per site, Tajima’s *D* (Tajima 1989) calculated in 10kb windows, and the number of singletons per site. Points represent the mean values, whilst confidence intervals represent the variance.

### Inferring the population history of common marmosets using a tailored ABC approach

The first step in demographic inference is to infer the number of populations implied by the empirical data. Based on cross-validation errors (CVE) produced by ADMIXTURE (Alexander et al. 2009), a single population was found to be most likely (*k* = 1, CVE = 0.57035; *k* = 2, CVE = 0.74668; *k* = 3, CVE = 0.97710; *k* = 4, CVE = 1.21181; *k* = 5, CVE = 1.28517; with *k* being the number of demes). We therefore inferred parameters for a single population demographic model in which the population may potentially experience both contraction and/or growth. The model contained four parameters: *N_ancestral_* (the ancestral diploid population size), *N_change_* (the proportionate instantaneous change in population size from *N_ancestral_*), *T_change_* (the time since the instantaneous size change in *N_ancestral_* generations), and *N_current_* (the population size at the time of sampling, with the size change between the instantaneous size change event and the population size at time of sampling occurring via exponential growth or decline). Parameters were initially drawn from a uniform distribution with ranges: 1,000 ≤ *N_ancestral_* ≤ 50,000; 0.01 ≤ *N_change_* ≤ 2; 0.01 ≤ *T_change_* ≤ 5; 1,000 ≤ *N_current_* ≤ 50,000, with the upper limit of *N_ancestral_* and *N_current_* increased to 80,000 following the first round of inference based on 1,000 draws from these priors. Given these parameter ranges, this model also allowed for no population size change (*i.e*., *N_change_* = 1 and *N_current_* = *N_ancestral_*), a single step-size change, and exclusive population growth or decline. Simulations for the ABC inference were performed in SLiM4 (Haller and Messer 2023), with 100 replicates for each parameter combination. Although any number of summary statistics might be used for inference with ABC, we found that *θ_w_*, Tajima’s *D*, and the number of singletons were the most informative summaries (further details of the ABC inference procedure are provided in the “Materials and Methods”).

Figure 3 depicts the posterior distributions for the four inferred parameters, the fit of the summary statistics for simulations across 50 ABC inference runs, as well as the inferred demographic model itself. The simulated demographic model utilizing the point estimates was found to fit the empirical data well, with the mean across simulation replicates capturing the mode of the empirical distribution. The values for these point estimates were *N_ancestral_* = 61,198; *N_change_* = 0.293; *T_change_* = 0.0287; and *N_current_* = 33,830, suggesting that the common marmoset underwent a major reduction in population size roughly 3,500 generations ago, before recovering to roughly half of its ancestral size. Notably, these parameter estimates are based on a mean mutation rate of 0.81 × 10^−8^ per base pair per generation inferred in a closely-related platyrrhine, owl monkeys (Thomas et al. 2018). Although a direct mutation rate estimate exists for common marmosets (0.43 × 10^−8^ per base pair per generation; Yang et al. 2021), this estimate was based on a single trio and the authors did not correct for the effects of chimerism. Thus, while the shape of the inferred demographic model would remain well-fitting, the parameter values themselves would vary depending on the mutation rate scaling (*e.g*., the inferred population sizes would be close to twice as large under the mutation rate estimated by Yang et al. [2021]).

**Figure 3:**
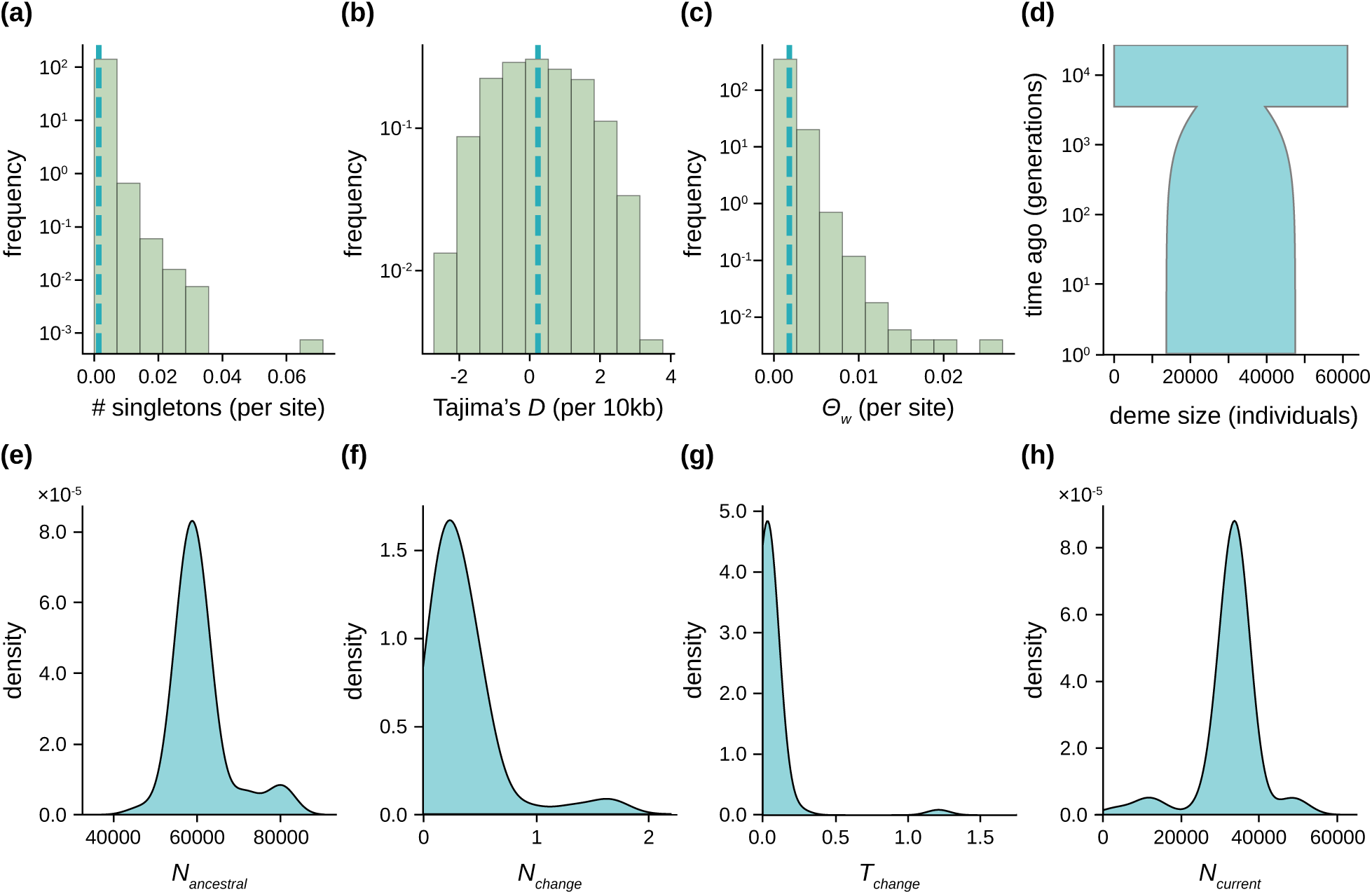
Results of the ABC demographic inference on the empirical marmoset data. **a-c)** Distribution of empirical summary statistics (the number of singletons per site, Tajima’s *D* [Tajima 1989] calculated in 10kb windows, and Watterson’s *θ* [*θ_w_*; Waterson 1975] per site, shown in green) and the mean values from simulations of mean parameter values from the posterior distributions (blue dashed line). **d)** Schematic of the demographic model with mean parameter values from the posterior distributions. **e-h)** Posterior distributions for four demographic model parameters: *N_ancestral_* (the ancestral diploid population size), *N_change_* (the proportionate instantaneous change in population size from *N_ancestral_*), *T_change_* (the time since the instantaneous size change, in *N_ancestral_* generations), and *N_current_* (the population size at time of sampling). Note that the differences in y-axis scaling owes to the differences in parameter range sizes.

### Inferring the population history of common marmosets using *δaδi*

We ran demographic inference across a range of 1D models with *δaδi* (Gutenkunst et al. 2009) in order to understand the extent of mis-inference when chimerism is unaccounted for (see “Materials and Methods”). The best-fitting model – based on the Akaike information criterion – was *δaδi*’s growth model, with inferred parameters of *Nu* = 0.402 and *T* = 0.270, where *Nu* is the ratio of ancient to contemporary population size and *T* is the time in the past at which growth began. Thus, these parameters represent an ancestral population size of 69,189, with the population undergoing an exponential decline that began 37,397 generations ago, reaching a current day population size of 27,832 diploid individuals (see Supplementary Figure S4 for a schematic of the best-fitting *δaδi* model, and Supplementary Figure S5 for the fit of the SFS between the inferred model and the empirical data). Although *δaδi* inferred similar ancestral and current day population sizes as our tailored ABC approach, the demographic models are different in shape, with no bottleneck event inferred by *δaδi*. Moreover, the ABC-inferred model includes a recovery post-bottleneck – albeit to a much-reduced size relative to the ancestral population size – whereas *δaδi* inferred a declining population. Crucially, *δaδi* inferred the decline as beginning over 10-fold generations earlier than the timing of our inferred bottleneck event. Our evaluation of neutral demographic estimators (see “Evaluating the impact of chimerism on the performance of common SFS-based estimators of population history”) demonstrated that the time of size change tends to be inflated by chimerism under models of population contraction (Supplementary Figures S2-S3), which likely explains the substantially older timing of the size change inferred by *δaδi* compared with the ABC model. Indeed, the posterior probability of such an old size change event is extremely low once accounting for chimerism (Figure 3g). Notably, and as shown in Supplementary Figure S5, there is nonetheless an appropriate fit between the SFS inferred by *δaδi* and that observed in the empirical data. These results therefore suggest that neglecting chimerism may result in a potentially mis-inferred but well-fitting demographic model.

## CONCLUSIONS

Although *C. jacchus* has been designated with the conservation status of least concern by the International Union for Conservation of Nature (Valença-Montenegro et al. 2021) – consistent with the relatively substantial population sizes here inferred – the species is of particular interest due to its frequent use in biomedical research (Vallender 2019), and because of the biologically peculiar phenomenon of chimerism. Assuming generation times in the common marmoset to be around 2 years (Tardif et al. 2003; Park et al. 2016; Schultz-Darken et al. 2016), the demographic model would place the inferred population collapse ∼7,000 years ago. This timing corresponds to a period during the Holocene epoch when forest habitats were eroded both by the expansion of human agriculture in north-eastern Brazil, and the expansion of savanna vegetation at the expense of arboreal vegetation, as evidenced in the carbon isotope record (Pessenda et al. 2004, 2010). Given that *C. jacchus* rely upon arboreal locomotion (Scheil and Souto 2017), this erosion of arboreal vegetation stands as a potential hypothesis for explaining the population reduction inferred in this study.

Overall, the well-fitting history inferred here based on a developed framework for modelling both twin-births and chimerism via forward-in-time simulation in an ABC framework, together with newly generated, high-quality, whole-genome, population-level genomic data, will prove valuable in future population genomic studies in this species. Specifically, this neutral baseline model stands as a necessary pre-requisite for further inference of both selective (*e.g*., the detection of selective sweeps and balancing selection) and neutral (*e.g*., the quantification of population-level recombination maps) processes (Johri et al. 2022b).

## MATERIALS AND METHODS

### Empirical data

#### Samples and sequencing

Genomic DNA (gDNA) from 15 common marmosets (*C. jacchus*) previously housed at the New England Primate Research Center was whole-genome sequenced to a target coverage of 35X per individual. All animals were maintained in accordance with the guidelines of the Harvard Medical School Standing Committee on Animals and the Guide for Care and Use of Laboratory Animals of the Institute of Laboratory Animal Resources, National Research Council. In brief, blood samples were collected during routine veterinary care under approved protocols. DNA was extracted using the FlexiGene kit (Qiagen, Valencia, CA) following manufacturer protocols. Prior to sequencing, the integrity of each gDNA sample was assessed using Agarose gel electrophoresis and the purity and concentration of samples were quantified using a NanoDrop Spectrophotometer and Qubit Fluorometer (ThermoFisher Scientific, Waltham, MA, USA), respectively. Afterward, a PCR-free library was prepared for each sample and paired-end sequenced (2 × 150 bp) on the DNBseq platform at the Beijing Genomics Institute (BGI Group, Shenzhen, China). Sample information and coverage statistics are provided in Supplementary Table S1.

#### Read mapping

Raw sequencing data was pre-processed using SOAPnuke v.1.5.6 (Chen et al. 2018) to remove adaptor sequences, contamination, and low-quality reads (using the following command line options: ‘ *-n* 0.01 *-l* 20 *-q* 0.3 *-A* 0.25 *--cutAdaptor -Q* 2 *-G --polyX --minLen* 150 ‘). Potentially remaining adapter sequences were marked using the Genome Analysis Toolkit (GATK) *MarkIlluminaAdapters* tool v.4.1.8.1 (van der Auwera and O’Connor 2020). Next, reads were mapped to the *C. jacchus* genome assembly of the Vertebrate Genomes Project (consisting of the maternal assembly for all autosomes and chromosome X and the paternal assembly for the Y chromosome; GenBank accession numbers: GCA_011078405.1 and GCA_011100535.1; Yang et al. 2021) using the Burrows Wheeler Aligner (BWA-MEM) v.0.7.17 (Li and Durbin 2009). To improve alignments, duplicates were marked using GATK *MarkDuplicates* v.4.2.6.1 prior to variant calling.

#### Variant calling, genotyping, and filtering

For each individual, germline variants were called from high-quality mappings (’ *--minimum-mapping-quality* 40 ‘) using the GATK *HaplotypeCaller* v.4.2.6.1 in base-pair resolution mode (’ *-ERC* BP_RESOLUTION ‘) to obtain calling information at each site of the genome. Thereby, the ‘ *--pcr-indel-model* ‘ was set to “NONE” as a PCR-free library protocol was followed during sequencing. Individual call sets were combined using GATK’s *CombineGVCFs* v.4.2.6.1 and jointly genotyped at all sites (’ *-all-sites* ‘) using GATK’s *GenotypeGVCFs* v.4.2.6.1. Next, the dataset was separated into autosomal, biallelic SNPs and monomorphic (*i.e*., invariant) sites genotyped in all individuals (AN = 30).

In the absence of a high-quality set of experimentally confirmed variants to train GATK’s Variant Quality Score Recalibration (VQSR) algorithm, variant sites were “hard” filtered using GATK’s *SelectVariants* and *VariantFiltration* tools v.4.2.6.1 following the GATK Best Practice recommendations (*i.e*., QD < 2.0, SOR > 3.0, FS > 60.0, RMSMappingQuality < 40.0, MappingQualityRankSumTest < −12.5, ReadPosRankSumTest < −8.0; van der Auwera and O’Connor 2020). In addition, as repetitive regions are prone to alignment errors and as extremely low or high read coverage is a frequent sign of sequencing and/or assembly issues (see discussion in Pfeifer 2017), SNPs located within repetitive elements as well as those supported by reads with less than half, or more than twice, the average individual autosomal depth of coverage were excluded from further analysis. To obtain information about the number of sites accessible to the study, invariant sites were, when applicable, subjected to the same filter criteria as variant sites.

#### Putatively neutral regions

To prevent biases in demographic inference due to positive selection, purifying selection, and/or background selection effects, variant and invariant sites were restricted to putatively neutral regions. Specifically, sites within 10kb of exons (based on 22,355 protein-coding genes annotated in the *C. jacchus* genome; Yang et al. 2021) as well as genomic regions conserved across the primate clade (Kuderna et al. 2024) were masked, resulting in a dataset of 1.4 million autosomal, biallelic SNPs with a transition-transversion ratio of 2.11 in the accessible genome (Supplementary Table S2). Summary statistics were calculated across 10kb windows with a 5kb step size using the Python implementation of libsequence v.1.8.3 (Thornton 2003), with means and standard deviations of the following statistics: the number of segregating sites (*S*), the number of singletons, *θ_w_* (Waterson 1975), and Tajima’s *D* (Tajima 1989).

### Evaluating the impact of chimerism on demographic inference

#### Simulations of chimerism in a non-WF framework

The non-WF framework in SLiM v.4.3 (Haller and Messer 2023) was used to simulate chimerism in dizygotic (fraternal) twins. In each generation, a random pair of individuals were joined as a monogamous breeding pair, with each subsequent reproductive event generating non-identical twins. At the end of each simulation, all individuals were sampled along with their pedigree IDs. For downstream inference, chimeric twins were subsampled post-simulation and their genotypes combined such that a mutation present in at least one of the twins was considered a SNP. For example, if one twin had a genotype of 0|1 and the other twin had a genotype of 1|1, the constructed chimeric genotype was considered to be 1|1.

#### Demographic model testing

In order to evaluate the impact of chimerism on demographic inference, simulations were performed under four population histories: (1) population equilibrium, (2) population expansion (instantaneous population size doubling), (3) population contraction (instantaneous population halving), and (4) severe population contraction (instantaneous population size reduction to 0.1*N*). For each population history, 100 replicates of a 1Mb region were simulated in a single population of 10,000 diploid individuals, with recombination rate of 1.0 × 10^−8^ per base pair per generation (Dumont and Payseur 2008) and a mutation rate of 2.5 × 10^−8^ per base pair per generation (Nachman and Crowell 2000) for testing purposes. After a burn-in of 14*N_ancestral_* generations (where *N_ancestral_* is the initial population size of 10,000), a size change occurred (when applicable) and samples were taken after an additional 0.01*N* generations. Afterward, ten chimeric and ten non-chimeric individuals were constructed as described in “Simulations of chimerism in a non-WF framework”.

Demographic inference was performed separately on chimeric and non-chimeric individuals using two commonly used demographic estimators: fastsimcoal2 (version fsc27; Excoffier et al. 2013) and *δaδi* (version 2.0.5; Gutenkunst et al. 2009). In the equilibrium model, a single population size parameter – the current population size (*N_current_*) – was inferred using fastsimcoal2 whereas in *δaδi, θ* was first estimated from the SFS and *N_current_* calculated from *θ*. In the population size change models, simulated spectra were fitted to the instantaneous size change (growth/decline) model in fastsimcoal2, which fits three parameters: the ancestral population size (*N_ancestral_*), the current population size (*N_current_*), and the time of change (*τ*). The parameter search ranges for *N_ancestral_* and *N_current_* were specified to be uniformly distributed between 10 and 100,000 individuals, whilst the range for *τ* was specified to be uniform between 10 and 10,000 generations. For size change inference in *δaδi,* the *two_epoch* model was used to fit two parameters: the current population size relative to the ancestral population size (*Nu_opt_*) and the time of change relative to the ancestral population size (*τ_opt_*), with *N_current_* once more calculated from *θ*. For each simulation replicate, 15 starting values between −2 and 2 were drawn for *Nu* and eight starting values between −2 and 2 were drawn for *τ*, both evenly distributed in log space, creating a total of 120 different starting parameterizations.

In fastsimcoal2, inference was conducted 100 times per simulation replicate, with 100 optimization cycles per run, and 500,000 coalescent simulations to approximate the expected SFS in each cycle. The best fit was that with the smallest difference between the maximum observed and the maximum estimated likelihood. In *δaδi*, the maximum number of iterations for the optimizer was set to 100 to facilitate convergence. The best fit for each simulation replicate was that with the lowest log-likelihood score.

### Inferring the population history of common marmosets

As the initial step in estimating the demographic history of the common marmoset, the extent of population structure was determined by ADMIXTURE v.1.3.0 (Alexander et al. 2009), using a range of *k* values from 1 to 5, where *k* is the number of demes. The number of demes and deme assignment of individuals was based on the value of *k* with the lowest CVE.

#### Demographic inference using ABC on empirical data

Following the approach described in “Simulations of chimerism in a non-WF framework”, 100 replicates of a 1Mb region were simulated in SLiM v.4.3 (Haller and Messer 2023). Thereby, mutation and recombination rate heterogeneity were modelled by sampling rates from a normal distribution with a mean of 0.81 × 10^−8^ per base pair per generation (the mutation rate inferred in a closely-related platyrrhine, owl monkeys; Thomas et al. 2018) and 1.0 × 10^−8^ per base pair per generation (the recombination rate observed in humans; Kong et al. 2002), respectively, and a standard deviation of a quarter of the mean.

For the ABC, parameters were drawn from a uniform distribution with ranges: 1,000 ≤ *N_ancestral_* ≤ 80,000; 0.01 ≤ *N_change_* ≤ 2; 0.01 ≤ *T_change_* ≤ 5; 1,000 ≤ *N_current_* ≤ 80,000, with the upper limit of *N_ancestral_* and *N_current_* increased to 80,000 following the first round of inference based on 1,000 draws from these priors. A further 100 draws were generated based on the posterior distribution generated using the “neural net” regression method with the default parameters provided by the R package “abc” (Csilléry et al. 2012). A 100-fold cross validation analysis was performed in order to determine the performance and accuracy of inference for tolerance values of 0.05, 0.08, and 0.1, with a tolerance of 0.08 identified as the most accurate. This value was employed for inference of final parameter values, meaning that 8% of all simulations were accepted by the ABC to estimate the posterior probability of parameter estimates. Inference was performed 50 times, with the mean of the weighted medians of the posterior estimates taken to determine point estimates of the inferred parameters. Finally, the demographic model was simulated in SLiM v.4.3 under these parameter values and plotted against the empirical distribution (Figure 3).

#### Demographic inference using δaδi on empirical data

For comparison with the ABC results, demographic inference on the empirical data was also performed using *δaδi* (Gutenkunst et al. 2009). Specifically, inference was performed on five 1D demographic models, supplied as part of the *δaδi* package: the standard neutral model (*SNM*), as well as *two_epoch*, *three_epoch*, *growth* and *bottlegrowth* models (see *δaδi* documentation for further details). Inference was performed 100 times, with 300 maximum iterations per run. Supplementary Table S3 lists the parameter ranges for each demographic model. The best-fitting model was determined by calculating the Akaike information criterion (Akaike 1974), which weights the likelihood of the model by the number of model parameters.

## ACKNOWLEDGEMENTS

We would like to thank the members of the Jensen and Pfeifer Labs for helpful comments and discussion. Library preparation and sequencing was performed at the Beijing Genomics Institute (BGI Group, Shenzhen, China). Computations were performed on the Sol supercomputer at Arizona State University (Jennewein et al. 2023).

---

**Supplementary Table S1.** Samples and coverage statistics. The 15 individuals (seven females and eight males) exhibited a genome-wide average coverage of 36.1X (range: 32.6-39.3).

| **sample** | **sex** | **coverage** |
| --- | --- | --- |
| CJac_01 | female | 39.0 |
| CJac_02 | male | 38.6 |
| CJac_03 | female | 39.3 |
| CJac_04 | female | 35.9 |
| CJac_05 | female | 34.6 |
| CJac_06 | male | 34.1 |
| CJac_07 | male | 33.9 |
| CJac_08 | male | 32.6 |
| CJac_09 | female | 33.4 |
| CJac_10 | female | 37.0 |
| CJac_11 | female | 33.6 |
| CJac_12 | male | 35.7 |
| CJac_13 | male | 39.3 |
| CJac_14 | male | 37.2 |
| CJac_15 | male | 37.2 |

**Supplementary Table S2.** Summary of variant and invariant datasets. 1.4 million autosomal, biallelic single nucleotide polymorphisms (SNPs) with a transition-transversion ratio (Ts/Tv) of 2.11 were discovered in the accessible genome of the 15 individuals included in this study.

| **chr** | **length** | **# SNPs** | **# invariant sites** | **Ts/Tv** |
| --- | --- | --- | --- | --- |
| 1 | 216,975,769 | 96,744 | 15,263,939 | 2.16 |
| 2 | 202,808,461 | 106,023 | 9,110,343 | 2.09 |
| 3 | 189,455,076 | 75,515 | 11,425,029 | 1.98 |
| 4 | 173,414,057 | 94,485 | 14,288,070 | 2.07 |
| 5 | 161,716,902 | 84,267 | 16,338,011 | 2.16 |
| 6 | 159,674,559 | 89,767 | 12,609,503 | 2.09 |
| 7 | 156,129,252 | 97,679 | 15,111,341 | 2.21 |
| 8 | 126,104,592 | 74,198 | 10,414,091 | 2.10 |
| 9 | 132,900,640 | 57,563 | 9,490,740 | 2.08 |
| 10 | 136,971,485 | 77,937 | 10,687,547 | 2.13 |
| 11 | 128,529,401 | 67,064 | 10,271,014 | 2.07 |
| 12 | 123,697,827 | 80,819 | 12,338,980 | 2.24 |
| 13 | 117,638,787 | 70,961 | 12,911,347 | 2.09 |
| 14 | 112,969,529 | 65,471 | 10,588,490 | 2.11 |
| 15 | 98,621,277 | 52,741 | 7,567,511 | 2.09 |
| 16 | 98,110,366 | 53,098 | 8,105,046 | 2.01 |
| 17 | 74,700,100 | 36,990 | 5,549,095 | 1.97 |
| 18 | 47,063,576 | 21,500 | 3,240,007 | 2.02 |
| 19 | 50,540,917 | 42,130 | 5,485,967 | 2.11 |
| 20 | 44,412,365 | 35,006 | 5,331,333 | 2.30 |
| 21 | 50,614,742 | 37,675 | 4,265,712 | 1.99 |
| 22 | 49,900,457 | 16,931 | 2,108,425 | 2.28 |
| **Σ or Ø** | **2,652,950,137** | **1,434,564** | **212,501,541** | **2.11** |

**Supplementary Table S3:** Parameter ranges for the demographic inference using *δaδi* on the empirical data.

| model | *Nu* |  | *T* |  | *NuB* |  | *NuF* |  | *TB* |  | *TF* |  |
| --- | --- | --- | --- | --- | --- | --- | --- | --- | --- | --- | --- | --- |
|  | lower | upper | lower | upper | lower | upper | lower | upper | lower | upper | lower | upper |
| *SNM* | - | - | - | - | - | - | - | - | - | - | - | - |
| *two_epoch* | 0.001 | 10 | 0.001 | 5 | - | - | - | - | - | - | - | - |
| *three_epoch* | - | - | - | - | 0.001 | 0.5 | 0.001 | 10 | 0.0001 | 0.05 | 0.0001 | 0.05 |
| *growth* | 0.001 | 10 | 0.001 | 5 | - | - | - | - | - | - | - | - |
| *bottlegrowth* | - | - | 0.001 | 5 | 0.001 | 0.5 | 0.001 | 10 | - | - | - | - |

**Supplementary Figure S1:**
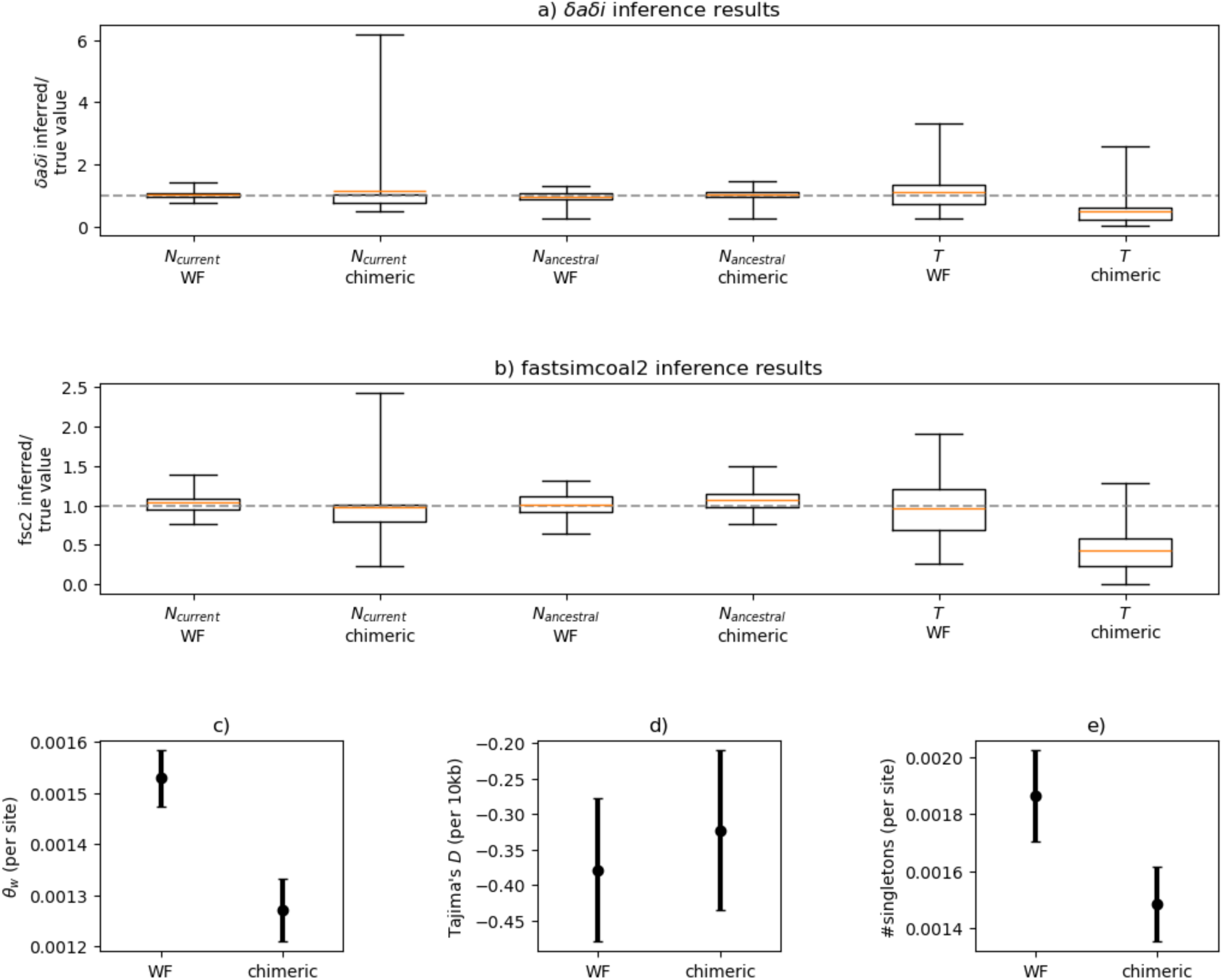
Demographic inference and summary statistics for simulations of a neutrally evolving population that has instantaneously doubled in size 0.01*N_ancestral_* generations ago (where *N_ancestral_* is the population size prior to the size change), across 100 simulation replicates, comparing Wright-Fisher (WF) and chimeric models. Demographic inference results from **a)** *δaδi* (Gutenkunst et al. 2009) and **b)** fastsimcoal2 (Excoffier et al. 2013). The y-axis represents the inferred value of the parameters relative to the true value of these parameters (with a value of 1 indicating that the inferred and true values are in agreement). Three parameters were inferred: *N_ancestral_*, *N_current_* (the population size at the time of sampling), and *T* (the time of size change). The orange line represents the mean inference value across 100 simulation replicates, with boxes representing 25 and 75 percentiles, and whiskers representing minimum and maximum values. **c-e)** Summary statistics, including Watterson’s *θ* (*θ_w_*; Waterson 1975) per site, Tajima’s *D* (Tajima 1989) calculated in 10kb windows, and the number of singletons per site. Points represent the mean values, whilst confidence intervals represent the variance.

**Supplementary Figure S2:**
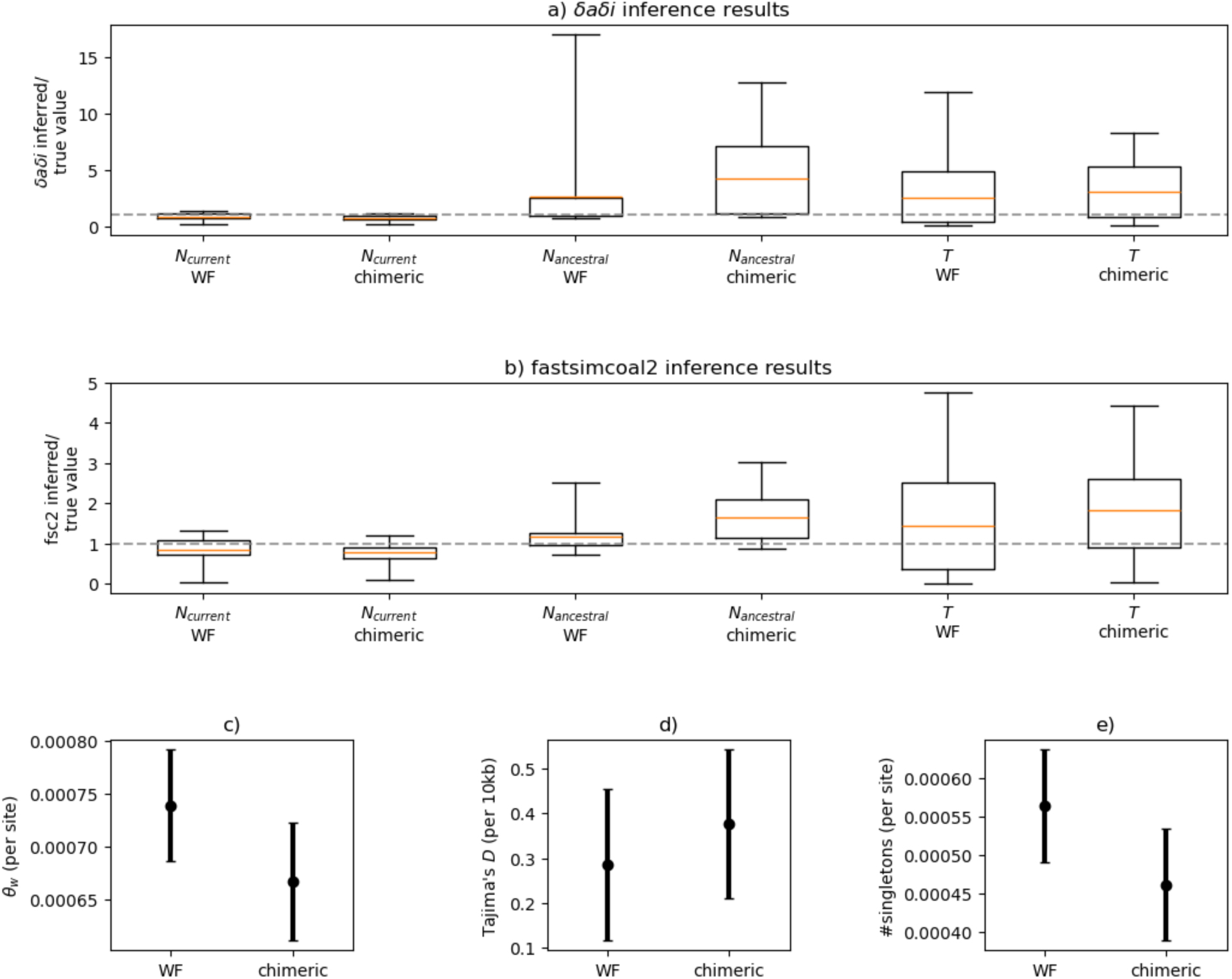
Demographic inference and summary statistics for simulations of a neutrally evolving population that has instantaneously halved in size 0.01*N_ancestral_* generations ago (where *N_ancestral_* is the population size prior to the size change), across 100 simulation replicates, comparing Wright-Fisher (WF) and chimeric models. Demographic inference results from **a)** *δaδi* (Gutenkunst et al. 2009) and **b)** fastsimcoal2 (Excoffier et al. 2013). The y-axis represents the inferred value of the parameters relative to the true value of these parameters (with a value of 1 indicating that the inferred and true values are in agreement). Three parameters were inferred: *N_ancestral_*, *N_current_* (the population size at the time of sampling), and *T* (the time of size change). The orange line represents the mean inference value across 100 simulation replicates, with boxes representing 25 and 75 percentiles, and whiskers representing minimum and maximum values. **c-e)** Summary statistics, including Watterson’s *θ* (*θ_w_*; Waterson 1975) per site, Tajima’s *D* (Tajima 1989) calculated in 10kb windows, and the number of singletons per site. Points represent the mean values, whilst confidence intervals represent the variance.

**Supplementary Figure S3:**
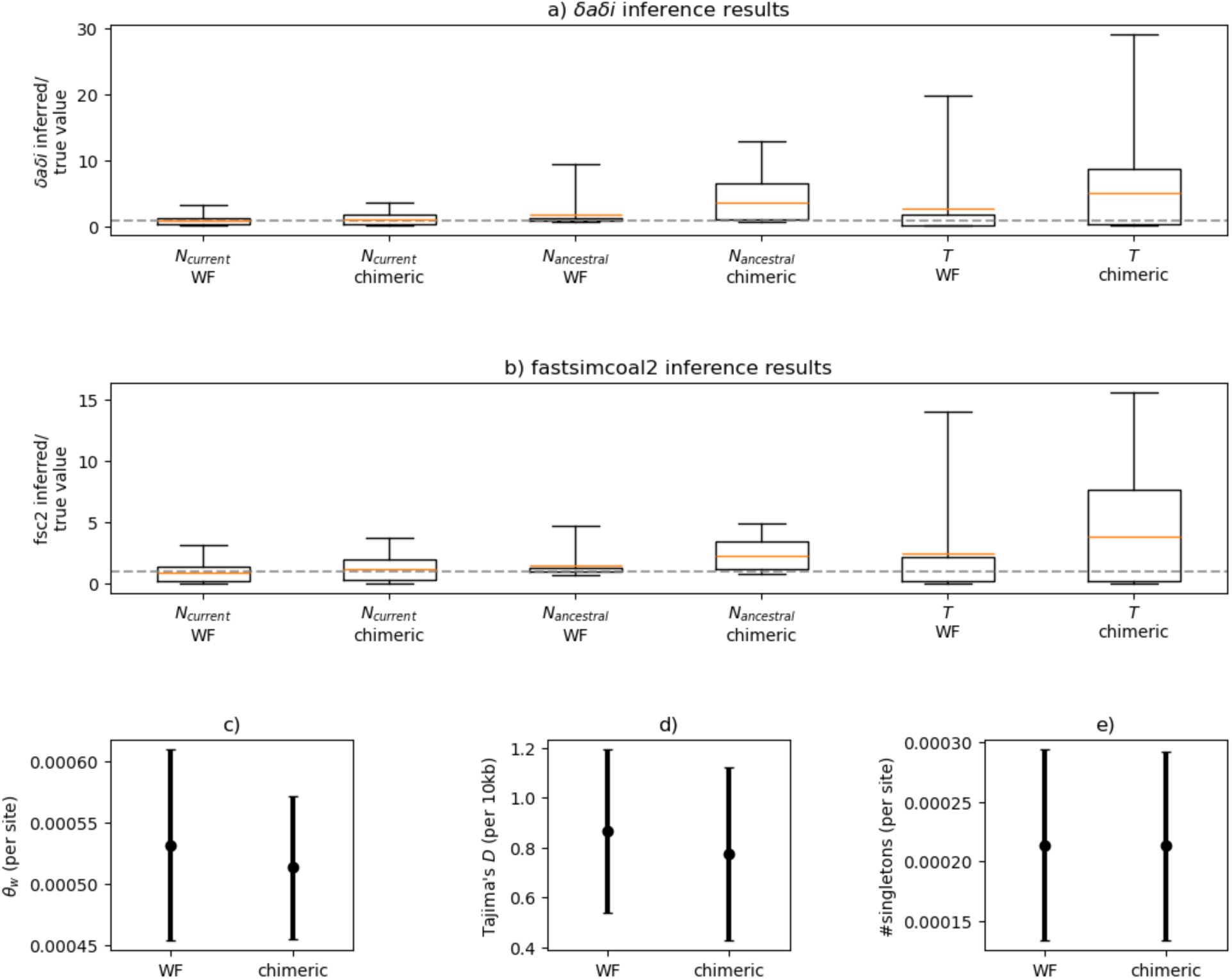
Demographic inference and summary statistics for simulations of a neutrally evolving population that has instantaneously decreased by 90% in size, 0.01*N_ancestral_* generations ago (where *N_ancestral_* is the population size prior to the size change), across 100 simulation replicates, comparing Wright-Fisher (WF) and chimeric models. Demographic inference results from **a)** *δaδi* (Gutenkunst et al. 2009) and **b)** fastsimcoal2 (Excoffier et al. 2013). The y-axis represents the inferred value of the parameters relative to the true value of these parameters (with a value of 1 indicating that the inferred and true values are in agreement). Three parameters were inferred: *N_ancestral_*, *N_current_* (the population size at the time of sampling), and *T* (the time of size change). The orange line represents the mean inference value across 100 simulation replicates, with boxes representing 25 and 75 percentiles, and whiskers representing minimum and maximum values. **c-e)** Summary statistics, including Watterson’s *θ* (*θ_w_*; Waterson 1975) per site, Tajima’s *D* (Tajima 1989) calculated in 10kb windows, and the number of singletons per site. Points represent the mean values, whilst confidence intervals represent the variance.

**Supplementary Figure S4:**
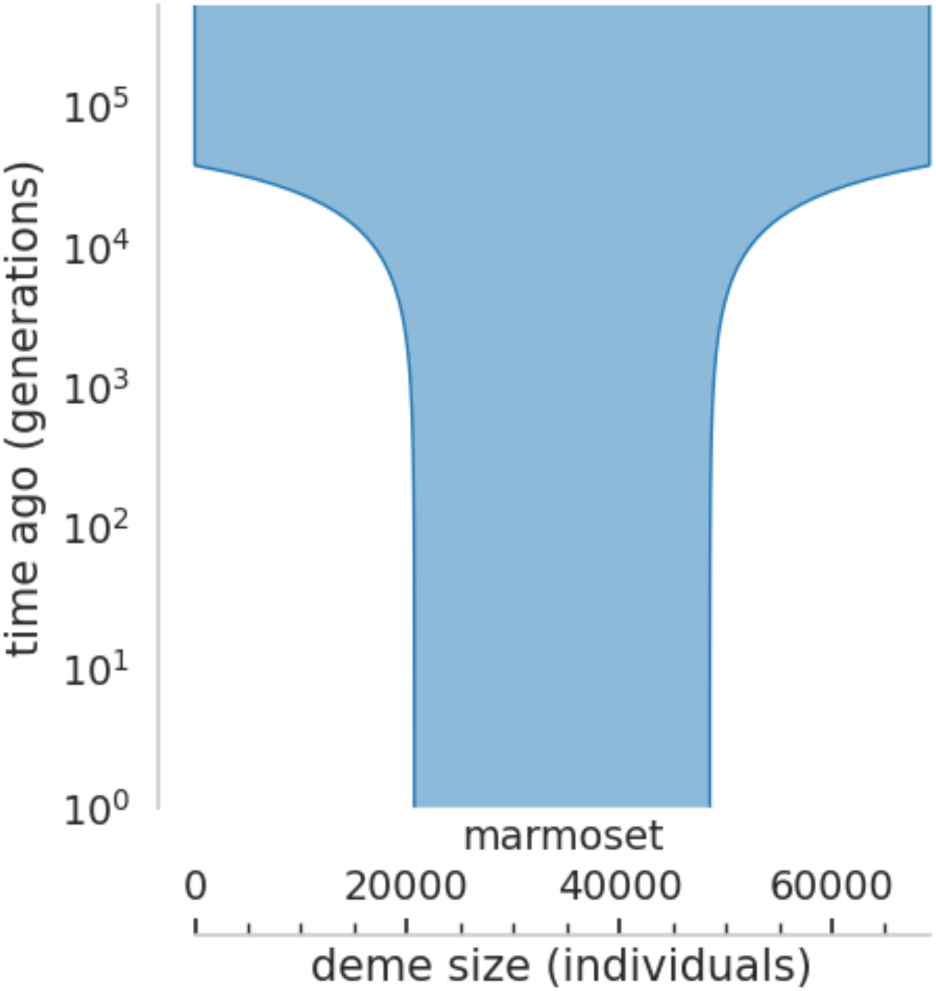
Schematic of the demographic model with the best-fitting parameters inferred from *δaδi*. Parameters for this model are *Nu* = 0.402 and *T* = 0.270, where *Nu* is the ratio of ancient to contemporary population size and *T* is the time in the past at which growth began.

**Supplementary Figure S5:**
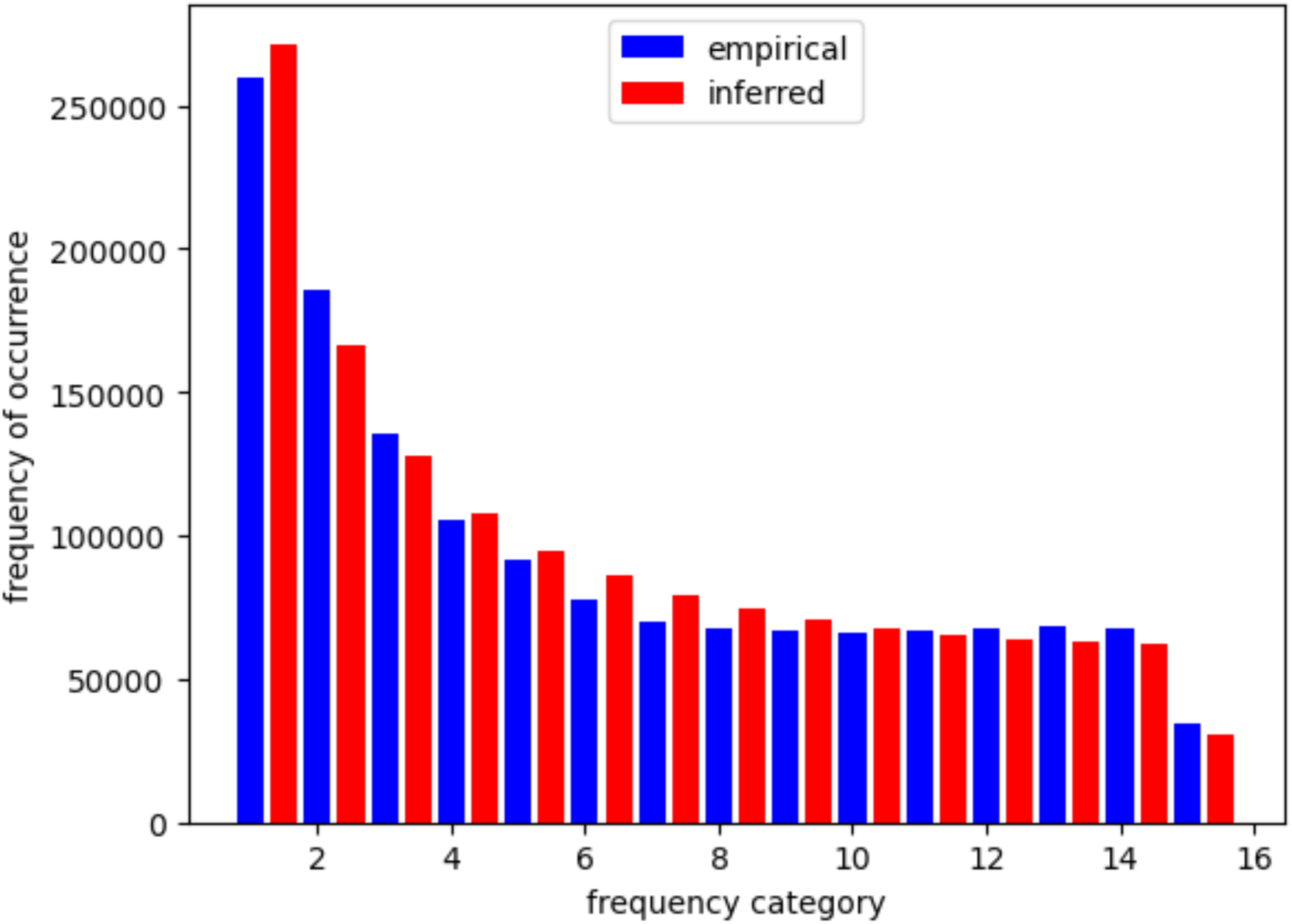
The site frequency spectrum for the best-fitting *δaδi* growth model (shown in red), compared to the empirical data (shown in blue). Parameters for this model are *Nu* = 0.402 and *T* = 0.270, where *Nu* is the ratio of ancient to contemporary population size and *T* is the time in the past at which growth began.

